# Does Zipf’s law of abbreviation shape birdsong?

**DOI:** 10.1101/2023.12.06.569773

**Authors:** R. Tucker Gilman, CD Durrant, Lucy Malpas, Rebecca N. Lewis

**Affiliations:** Department of Earth and Environmental Sciences; Faculty of Science and Engineering; University of Manchester; Manchester, UK; School of Biological Sciences; Faculty of Biology, Medicine, and Health; University of Manchester; Manchester, UK; Department of Natural Sciences; Manchester Metropolitan University; Manchester, UK; Chester Zoo; Upton-by-Chester, Chester, UK

## Abstract

Zipf’s law of abbreviation predicts that in human languages, words that are used more frequently will be shorter than words that are used less frequently. This has been attributed to the principle of least effort – communication is more efficient when words that are used more frequently are easier to produce. Zipf’s law of abbreviation appears to hold for all human languages, and recently attention has turned to whether it also holds for animal communication. In birdsong, which has been used as a model for human language learning and development, researchers have focused on whether more frequently used notes or phrases are shorter than those that are less frequently used. Because birdsong can be highly stereotyped, have high interindividual variation, and have phrase repertoires that are small relative to human language lexicons, studying Zipf’s law of abbreviation in birdsong presents challenges that do not arise when studying human languages. In this paper, we describe a new method for assessing evidence for Zipf’s law of abbreviation in birdsong, and we introduce the R package ZLAvian to implement this analysis. We used ZLAvian to study Zipf’s law of abbreviation in the songs of 11 bird populations archived in the open-access repository Bird-DB. We did not find strong evidence for Zipf’s law of abbreviation in any population when studied alone, but we found weak trends consistent with Zipf’s law of abbreviation in 10 of the 11 populations. Across all populations, the negative correlation between phrase length and frequency of use was several times weaker than the negative correlation between word length and frequency of use in human languages. This suggests that the mechanisms that underlie this correlation may be different in birdsong and human language.

## Introduction

Over the past three decades, birdsong has gained currency as a tractable model for studying how language develops and is transmitted in humans [1–6]. This has been due in part to the discovery of biological similarities between birdsong and human speech, including analogies in learning patterns [1, 4, 6], brain mechanisms [7], and regulatory genetics [1, 8]. Birdsong is also amenable to experimentation that might be impractical or unethical in humans [1]. These attributes have made birdsong particularly appealing as a model system for studying human speech pathologies [9–11]. The growing importance of birdsong as a model of human language necessitates a clearer understanding of how birdsong and human language are similar and how they differ, so we can better understand both the potential applications and the limitations of the model [12].

Zipf’s law of abbreviation (ZLA) states that, in human languages, words that are used more frequently tend to be shorter than words that are used less frequently. This has been attributed to the principle of least effort. If an idea must be conveyed frequently, users will find or create shorter words to convey that idea, thus making communication more efficient [13, 14]. If users must convey an idea only infrequently, then they can invest effort in longer words to ensure that the idea is communicated clearly [14]. Evidence supporting ZLA has been found in each of the nearly 1,000 human languages where it has been sought [15], and the law applies to both spoken language [16, 17] and written characters [18, 19]. Researchers have reported mixed support for ZLA in the vocal communication of other animals [20], including primates [17, 21–25], cetaceans [26], bats [27], and hyraxes [28]. Relatively few studies have looked for patterns consistent with ZLA in birds. More than 30 years ago, Hailman and colleagues [29] reported that in black-capped chickadees (*Parus atricapillus*) shorter bouts of calls were more frequent than longer bouts of calls, but they found no evidence that shorter call types were more frequent than longer call types. This has been cited as an example of ZLA in birds [22, 24, 30, 31], but it is not clear that the pattern Hailman and colleagues [29] reported should emerge due to the mechanism Zipf [13] proposed. Birdsong or calls can be segmented into notes (continuous sounds separated by periods of silence), phrases (short series of notes that frequently or always appear together), calls or songs (series of notes or phrases separated by longer periods of silence), and bouts (series of often similar calls or songs separated by even longer periods of silence) [6, 12, 29]. If notes or phrases are analogous to words (which is debated [32]), then calls and bouts may be analogous to sentences and orations, respectively. ZLA posits a relationship between the frequency and length of words, and it is not clear that the same relationship should emerge at these higher levels. Indeed, a simple process in which birds begin bouts of calls and then decide independently after each call whether to stop or continue could produce patterns similar to those Hailman and colleagues [29] reported. In 2013, Ferrer-i-Cancho and Hernández-Fernández [17] found no evidence for ZLA in the calls of the common raven (*Corvus corax*) in data collected by Connor [33]. In 2020, Favaro and colleagues [34] reported that shorter note types appear more frequently than longer note types in the calls of captive African penguins (*Spheniscus demersus*). However, the study population used only three note types, so the perfect negative concordance between note duration and frequency of use that the authors observed could easily have arisen by chance. More recently, Lewis and colleagues [35] found no evidence for ZLA in the songs of a domesticated population of Java sparrows (*Padda oryzivora*). Thus, whether there are patterns consistent with ZLA in bird vocalizations remains an open question.

Given the currently weak evidence for ZLA in bird vocalizations, one might reasonably ask whether we should expect to see ZLA in birds at all. In human languages, words have lexical meanings, and those meanings can be independent of the length of the word. For example, we can shorten “television” to “TV” or “telly” and the meaning does not change. In birdsong, the sound of a note may determine its value to the listener [12]. For example, in some species, females appear to interpret specific note types as indicators of male quality, perhaps because those note types are difficult to produce [36–39]. If a male produced shorter or longer versions of those note types, then the information conveyed to females about his quality might change. Thus, replacing long note types with short ones might not make communication more efficient but rather prevent accurate communication among birds.

### Challenges to studying Zipf’s law of abbreviation in birdsong

Assessing the evidence for ZLA in birdsong presents several challenges that we do not encounter when studying ZLA in humans. First, relative to the number of words in human languages, the number of note types used by most bird populations is small. A small number of note types makes it more difficult to detect a significant concordance between note type frequency and duration [15, 17]. If the number of note types is very small, as in the calls of African penguins [34], then even a perfect concordance between the frequency and duration of note types may provide only weak evidence for or against ZLA. No amount of additional study can resolve this problem. Thus, we may never be able to say with confidence that ZLA operates in particular populations. Instead, researchers interested in ZLA in birdsong may need to assess large numbers of populations and draw conclusions based on the full body of evidence.

A second challenge stems from the fact that different birds in the same population can have very different note type repertoires [40]. In humans, individuals in a population that shares a language are likely to use similar sets of words with similar frequencies. Thus, it may be reasonable to study ZLA at the population level. Researchers can select representative multiauthor texts and assess the concordance between the frequency and duration for each word using simple rank correlations [15]. In contrast, in many bird species, individuals in the same population use different and sometimes non-overlapping sets of note types [40]. This makes it difficult to adequately sample the use of note types in those populations. The problem is compounded by the fact that, in at least some species, song durations themselves appear to be constrained [41, 42]. Birds that use longer note types sing fewer notes in each song. In such species, if birds that sing shorter note types are at least as common as birds that sing longer note types, then we might see patterns consistent with ZLA at the population level even if no individual bird uses short note types more frequently than it uses long ones. However, such a pattern would not provide evidence for the principle of least effort proposed to underlie ZLA.

Because the principle of least effort suggests that individuals should use shorter types (ie, words or notes) more frequently than longer ones, we might wish to look for ZLA at the level of individuals rather than populations. That is, if we choose a random individual from a population, are we likely to find that this individual uses shorter note types more frequently than longer ones? However, this question is made difficult by the fact that songs produced by individual birds in the same population may not be independent. In many species, songs are highly stereotyped and birds learn their songs from others [40, 43, 44]. If we find that two birds have note use consistent with ZLA, then the pattern may have arisen independently in each bird, or it may have arisen only once and both birds may have learned it from the same source. The second case is weaker evidence for ZLA. Thus, any attempt to study ZLA at the level of individuals must adequately account for the potential non-independence of individuals’ songs.

Finally, perhaps the biggest challenge to studying ZLA in birdsong arises from the inherent difficulty of classifying notes. In human languages, especially in the written form, we can usually agree on whether two units represent the same word or different words [13, 15, 45]. In birdsong, determining whether two notes belong to the same note type is less straightforward. Notes are usually assigned to types by expert inspection of spectrograms [40, 46] or sometimes by computational clustering [47–49]. Both methods are highly repeatable [40, 49]. However, high repeatability does not ensure that the assigned note types match the intent of the birds that produced those notes. Different birds may produce notes that are very similar but are nonetheless objectively distinguishable among individuals (eg, because they have slightly different peak frequencies or durations [40]). Should we assign these notes to the same or different types? Similarly, individual birds may produce objectively distinguishable versions of similar notes at different points in their song. In general, we cannot know whether the bird intends to produce slightly different notes, or whether it is attempting to produce the same note each time but its performance is constrained by the position of the note in the song. It is not clear that we can resolve this problem empirically. We could ask whether listening birds can distinguish between notes, but the ability of other birds to distinguish between notes does not necessarily indicate the intent of the producer. By analogy, I might attempt to imitate a word pronounced by my colleague, but listeners may still be able to distinguish my attempt from theirs, even though I am attempting to produce the same sound. In some study populations (eg [40]), we may know which birds learned their songs from which others, and we may be able to use analogies in song structure to infer that similar-sounding notes are attempts to produce the same note type. However, for most populations, we do not know which birds learned from which others, and inferring the intent behind individual notes may be fundamentally beyond our grasp. In cases where it is difficult to decide whether notes should be assigned to the same or different types, it is likely that the notes will have similar characteristics, and therefore the decision to split or merge types may have little effect on the durations of the note types. However, the decision to split or merge types will necessarily affect the frequencies with which those types appear, and so may affect our inferences about ZLA.

### A method for assessing Zipf’s law of abbreviation in birdsong

With these challenges in mind, Lewis and colleagues [35] developed a method for assessing ZLA in bird populations. In this paper, we introduce the R package ZLAvian to implement that method. The analysis requires birdsong data in which notes have been assigned to types, the duration of each note has been measured, and each note can be attributed to an individual bird. We compute the mean logged duration of each note type as produced by each bird in the sample, and we count the number of times that each bird produced each note type. Then, we compute the concordance between the duration and the frequency with which each note type is produced (ie, Kendall’s τ) within birds. This results in one value of τ for each bird. We compute 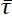, the population mean value of τ with each bird weighted by the inverse variance of its τ. Weighting by the inverse variance of τ accounts for the fact that we can estimate τ more accurately in birds that have larger note type repertoires [50]. Our 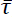 serves as a test statistic, but also has a clear biological interpretation. If we were to randomly select a bird from the study population and then randomly select two note types as produced by that bird, then 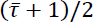 is the probability that the longer note type would appear more frequently. This makes 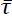 a useful and intuitive metric for comparing the observed strength of ZLA across populations.

Next, we computed a null distribution for 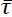. To do this, we first computed the expected logged duration for each note type in the population. We obtained the expected logged duration for each note type from a model of the observed logged durations for that note type with a random effect of the bird that produced each individual note. This method accords more weight to birds that produce the note types more frequently, because we can better estimate the note type durations in those birds. Then, we permuted the expected logged durations among the note types at the population level. Thus, if a note type was assigned a particular duration by permutation, then it was assigned that same duration in all birds that produced that note type. This permutation results in a set of population mean logged durations for note types that we might see under the null hypothesis that note type durations and frequencies of use are independent, but it maintains the observed distribution of note type frequencies within birds in the population. This accounts for the fact that birds may learn songs from other birds.

In nature, individual birds may produce the same note type in slightly different ways, and therefore the mean logged duration of each note type as produced by each bird will differ from the mean logged duration of that note type in the population. As a result, the rank order of note type durations can differ among birds. How each bird produces each note type may be learned from other birds, and durations may not be independent among the note types a bird produces. For example, a bird that produces longer versions of one note may be more likely to produce longer versions of other notes. We want to account for differences in note duration among birds under maximally conservative assumptions about how note type versions are learned. To do this, we first computed the deviation of each bird’s mean logged note duration from the population mean for each note type that the bird produced. Then we either added or subtracted these deviations from the permuted population mean logged durations for each note as produced by each bird. We treated the set of deviations as a block – that is, if we added the deviation to one note type as produced by one bird in the data set, then we added the deviations to all note types as produced by all birds in the data set. This allows the rank order of note durations to differ among bird as in the observed data, but maintains the structure of deviations in note durations within and among birds. We then computed 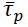, the analog of 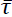 for the permuted data set. The null distribution of 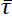 is the set of 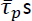 computed for every possible permutation of the mean logged note durations with the positive or negative deviation structures added. Except when the number of note types is very small, we estimated the null distribution from a randomly chosen subset of the possible permutations. The p-value for the hypothesis that ZLA operates in the study population is the proportion of the null distribution in which 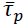 is equal to or smaller than 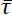. Lewis and colleagues [35] developed this method to study the concordance between note type durations and frequencies of use in birdsong, but the method is transferable to other taxa and to other measures production effort (eg, bandwidth, concavity, excursion [51]).

The method proposed by Lewis and colleagues [35] assesses evidence for ZLA within individuals while accounting for non-independence among individuals due to song learning or stereotypy. However, it does not correct for flaws in the assignment of note types. Such flaws result in errors in the data, and in general statistical methods cannot correct errors in the data. However, we can assess how different kinds of note type misclassifications will affect our inferences (see supplementary information). If we incorrectly merge note types (ie, assign notes to the same type when they should belong to different types), we will overestimate the variance of the null distribution, and our inference will be conservative. If we incorrectly split note types, we will underestimate the variance of the null distribution, and our inference will be anticonservative. A more problematic flaw arises if we systematically misclassify note types. One plausible example is that we might be more likely to split longer note types, or merge shorter note types, simply because longer note types give us more opportunity to identify potential differences among notes. In this case, we would systematically bias 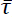 downwards, and we might infer ZLA in our data not because of how birds use note types but rather because of how we perceive the note types they use. There is no safeguard against this kind of misclassification, and attempts to assess ZLA in birdsong must be interpreted in light of this limitation.

### Motivation for the current study

Because of the potential for humans to systematically misclassify notes in birdsong, data from simple surveys of birdsong in populations may never conclusively demonstrate the existence of ZLA. Nonetheless, we believe such data is worth analysing, and we encourage authors to do so when they have appropriate data available. If we find little evidence for ZLA in birdsong, or if we find many populations in which the duration and the frequency of use of notes are positively correlated, then we might conclude that ZLA does not operate in birdsong, or at least that it is not universal in birdsong, as it appears to be in human language [15]. If we find widespread evidence for ZLA, then researchers may consider more focused studies to identify the potential mechanisms that produce ZLA. If we find evidence for ZLA in some bird populations but not others, then researchers can begin to study other ways in which these populations differ.

With these goals in mind, we assessed the evidence for ZLA in seven species of songbirds in an open access repository of annotated birdsong [52]. We conducted our analysis using the package ZLAvian (https://CRAN.R-project.org/package=ZLAvian) in R [53]. ZLAvian will facilitate similar analyses by other researchers who have or can obtain annotated birdsong data. Our results will contribute to a deeper understanding of the similarities and differences between birdsong and human language.

## Methods

We downloaded 660 annotations of birdsong representing seven different bird species (California thrasher, *Toxostoma redivivum*; redthroat, *Pyrrholaemus brunneus*; black-headed grosbeak, *Pheucticus melanocephalus*; sage thrasher, *Oreoscoptes montanus*; Cassin’s vireo, *Vireo cassinii*; western tanager, *Piranga ludoviciana*; gray shrikethrush, *Colluricincla harmonica*) from the open access repository Bird-DB [52] on 18 April 2022. Annotations on Bird-DB can include one or more songs, but all songs in the same annotation are from the same bird. The phrases in each annotation have been classified into types, and the starting and ending times for each phrase are reported. We used the reported starting and ending times to compute the phrase durations. The recordings represented in Bird-DB were collected on different days and at different locations, and we assumed that each annotation represents a different bird. Most phrases in Bird-DB are monosyllabic, and thus correspond to individual notes. A small number of phrases consist of short sequences of notes that typically appear together. Such polysyllabic phrases are often analysed as single units, and we follow this convention in the body of this paper. Our results are qualitatively similar if we divide the polysyllabic phrases into individual notes and use notes rather than phrases as the primary unit of analysis (see supplementary information).

We cleaned the annotations downloaded from Bird-DB prior to analysis. Phrase types in Bird-DB are identified by two- or three-letter strings. We excluded any phrase with a type identifier that includes non-alphabetic characters, or that comprises fewer than two or more than three characters. These are likely to be data entry errors, and we cannot confidently assign these phrases to types. We also excluded annotations that include only one repeated monosyllabic phrase type. These annotations may represent alarm calls, and alarm calls may adhere to different rules than other calls or songs. Assessing concordance within annotations requires at least two phrase types, so no information about concordance was lost by excluding annotations that consisted of only one phrase type.

In some cases, songs from the same species on Bird-DB were annotated under different classification systems. We cannot analyse these songs together, because we do not know which phrases in one classification system correspond to which phrases in the other. If we treat phrases from different classification systems as different when they are in fact the same, we will overestimate the number of phrases in the species’ repertoire and underestimate the variances of null distributions, and the p-values we obtain when testing for ZLA in that species will be anticonservative. Therefore, when songs from the same species were annotated using different classification systems, we treated annotations classified by each system as different populations. Tests conducted on multiple populations from the same species cannot be regarded as independent, because populations may share phrases and song structures. Nonetheless, tests that produce similar results using different populations can provide corroborating evidence for or against the presence of ZLA in that species. For each population represented in Bird-DB, we report the number of annotations studied, the total number of phrase types across all annotations, the mean number of phrases and phrase types that appear in each annotation, the mean Shannon diversity of phrase types in each repertoire, the value and statistical significance of the concordance between phrase type duration and frequency of use at the population level, and the mean value and statistical significance of the concordance between phrase type duration and frequency of use by individuals in the population (ie, 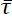). All of these measures are computed by the package ZLAvian.

If we find no evidence for ZLA in the birdsongs we studied, it may be because birdsong does not conform to ZLA. Alternatively, it may be because ZLA produces only weak concordances in birdsong, and the repertoire sizes of the individual populations we studied were too small to assign significance to those concordances. Therefore, we conducted a second analysis in which we compared the concordances we observed in birdsong to those in human languages. To represent ZLA in birdsong, we chose the concordance from the population of each species that used the largest number of phrases. We chose just one population per species because concordances in populations of the same species may not be independent. Then, following [15], we measured the length in characters and the frequency of use of words in 462 translations of the Universal Declaration of Human Rights obtained from https://unicode.org/udhr/index.html. We computed the concordance between the length and frequency of use of words in each translation. We compared the concordances we observed in birdsong to those we found in human languages with a t-test using Welch’s correction for unequal variance in the two groups. This analysis tells us not whether ZLA operates in birdsong, but instead whether ZLA differs in birdsong and in human language.

## Results

We assessed the evidence for ZLA in 11 populations from 7 bird species (table 1). Each population was represented by 2 to 296 annotations (mean 51.0, median 13, sd 87.2). The number of phrase types per population ranged from 9 to 748 (mean 188, median 114, sd 219) and the number of phrase types per annotation in the populations ranged from 2.8 to 89.5 (mean 30.0, median 24.9, sd 23.8).

Four of 11 concordances at the population level were significant (α = 0.10). Two of these were negative (ie, consistent with ZLA; one in California thrashers and one in western tanagers) and two were positive (ie, contrary to ZLA; in black-headed grosbeaks and Cassin’s vireos). Only one of the mean individual concordances was significantly different from the null expectation (consistent with ZLA in western tanagers) but 10 of the 11 were negative.

The mean of the mean individual concordances across the seven populations with the largest repertoires in our study was -0.066±0.028. The mean concordance we observed in human languages was -0.212±0.002. Thus, concordances were more negative in human languages than in birdsong (p = 0.002; figures 1, S3.1).

**Table 1.** Summary statistics and concordances in the songs of 11 populations of 7 bird species archived on Bird-DB. Concordances in bold are significantly different from the null expectation (α = 0.10). P-values less than 0.05 indicate patterns consistent with ZLA (black), and p-values greater than 0.95 indicate patterns contrary to ZLA (red).

| species | records studied | total phrase types | phrases record<sup>-1</sup> | phrase types record<sup>-1</sup> | Shannon diversity | concordance (population) | mean concordance (individual) |
| --- | --- | --- | --- | --- | --- | --- | --- |
| California thrasher | 89 | 748 | 146 | 14.4 | 2.20 | **-0.079 (p < 0.001)** | 0.022 (p = 0.820) |
|  | 7 | 181 | 412 | 57.4 | 3.62 | -0.073 (p = 0.079) | -0.063 (p = 0.095) |
| redthroat | 7 | 56 | 176 | 14.9 | 2.15 | -0.054 (p = 0.285) | -0.035 (p = 0.347) |
| black-headed grosbeak | 83 | 451 | 153 | 27.5 | 2.92 | -0.009 (p = 0.388) | -0.046 (p = 0.064) |
|  | 16 | 107 | 109 | 25.9 | 2.93 | **0.147 (p = 0.985)** | -0.036 (p = 0.298) |
| sage thrasher | 2 | 147 | 234 | 89.5 | 4.13 | -0.049 (p = 0.209) | -0.040 (p = 0.259) |
| Cassin's vireo | 13 | 68 | 87.7 | 26.5 | 2.98 | **0.161 (p = 0.970)** | -0.063 (p = 0.211) |
|  | 296 | 134 | 119 | 21.5 | 2.54 | -0.026 (p = 0.326) | -0.028 (p = 0.197) |
|  | 41 | 114 | 94.6 | 26.6 | 2.85 | 0.083 (p = 0.903) | -0.032 (p = 0.219) |
| western tanager | 3 | 56 | 129 | 22.7 | 2.30 | **-0.193 (p = 0.023)** | **-0.170 (p = 0.038)** |
| grey shrike-thrush | 4 | 9 | 13.3 | 2.8 | 0.76 | -0.032 (p = 0.455) | -0.166 (p = 0.311) |

**Figure 1.**
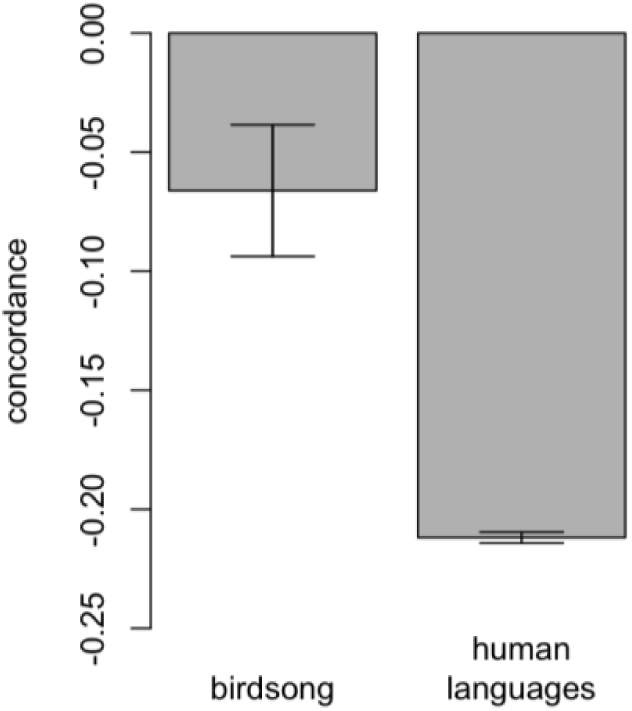
Mean concordance between phrase duration and frequency of use in the songs of seven bird species, and between word length and frequency of use in samples from 462 human languages. Error bars show standard errors. Concordances are more negative in human languages than in birdsong (p = 0.002).

## Discussion

Zipf’s law of abbreviation predicts a negative concordance between phrase duration and frequency of use within individuals in a population [13]. We found statistically significant evidence consistent with ZLA in only one of the 11 populations we studied. However, in 10 of 11 populations, the best estimates for the mean individual concordance were negative. Similar nonsignificant trends have been reported in the songs of Java sparrows [35] and in call repertoires at the population level in common ravens [17] and African penguins [34]. Taken together, this evidence is consistent with a weak effect of ZLA in bird vocalisations that is difficult to detect when phrase or note repertoires are small. It may be necessary to assess ZLA in many different bird species before we can draw clear conclusions about its existence or strength in birdsong generally. The results we present here are a first step towards this goal.

The birdsong phrases we analysed in our study were assigned to types by humans. We cannot know whether humans perceive phrases in the same way that birds do. When birds produce longer phrases, there may be more opportunities for error than when they produce shorter phrases. If a bird attempts to produce the same phrase type several times, if some of those attempts include errors, and if researchers interpret phrases with and without errors as different types, then we may systematically overestimate the number of long phrase types and underestimate the frequency with which each long phrase type is used in that bird’s repertoire. If researchers are less able to distinguish differences between phrase types when those phrase types are short, and if this leads researchers to merge phrase types that are different for birds, then we may systematically underestimate the number of short phrase types and overestimate the frequency of each short phrase type in birds’ repertoires. In either case, our tests for ZLA will be anticonservative. This cannot explain why we failed to find strong evidence of ZLA in the populations we studied, but it could explain or partly explain why we found trends towards negative concordances in most populations.

We assessed the concordances between phrase duration and frequency of use in populations and also within individuals. Concordances at the population level sometimes differed qualitatively from those at the individual level. Such patterns can arise if birds have different repertoires, and if some repertoires are more common or if birds with some repertoires are recorded more frequently than others. If individuals develop shorter versions of phrases they use more frequently, then we should expect concordances between phrase duration and frequency of use to arise within individuals. Many researchers have studied ZLA in animals at the population level [17, 20–28, 34, 51], but our results underscore the importance of verifying these patterns at the individual level.

The negative concordance between phrase duration and frequency of use that we observed in birdsong is several times weaker than the negative correlation between word length and frequency of use that we measured in human languages. This may indicate that birdsong and human language follow different organising principles, and suggest limitations in the value of birdsong as a model for human language learning or processing. One reason that ZLA may impact human languages more strongly than birdsong is that the nature of the tokens (ie, words in human languages, notes or phrases in birdsong) differs in the two systems. In human languages, words have lexical meanings or functions. By developing shorter versions of words they use more frequently, users can communicate more efficiently [13]. In birdsong, notes or phrases may not have meanings independent of the note or phrase itself [12]. In the context of courtship or territory defence, the primary function of birdsong may be to advertise the quality of the singer, and notes or phrases that are more difficult to produce may indicate higher quality [39, 54, 55]. If this is true, then it may be impossible to shorten the duration of note types without also changing the message conveyed to listeners. This would disable the mechanism that underlies ZLA in human languages. In some animals where patterns consistent with ZLA have been identified, the tokens studied are thought to have semantic meanings [21, 56], making these communication systems more similar than birdsong to human language.

In birds, alarm calls may offer an appealing context for studying ZLA. In some bird species, the phrases that make up alarm calls have specific meanings. For example, different phrases can indicate different predator types [57–59]. These phrases can differ among populations and change within populations over time [60]. This may allow ZLA to shape alarm calls more strongly than it shapes song. Studies of the concordance between phrase duration and frequency of use in alarm calls in bird populations under different predation pressures may reward effort.

Identifying patterns consistent with ZLA in birdsong, and quantifying those patterns if they exist, may require studying the songs of many different bird populations and species. This requires songs that can be attributed to individual animals, where notes or phrases have been classified to type, and where the durations of notes have been measured. This data already exists for many species, and automated note or phrase classification (eg, [47, 48]) may make such data easier to collect in the future. The ZLAvian package we introduce here will allow researchers who collect or maintain such data sets to quickly and easily test for evidence of ZLA. Thus, our work offers the opportunity to expand our understanding of the similarities and differences between birdsong and human language.

## Acknowledgments

The authors thank Patrycja Strycharczuk for advice and discussion.

## Supplementary information

Appendix 1: Robustness analysis showing how the inferred relationship between phrase type duration and frequency of use depends on plausible types of phrase classification errors.

Appendix 2: Robustness analysis showing that qualitative results are similar if we treat phrases as catalogued on Bird-DB or individual notes as tokens when assessing ZLA.

Appendix 3: Supplementary figure showing the within-individual concordances between phrase type duration and frequency of use in our study populations, and for comparison, the concordance between word duration and frequency of use in the first 10 psalms in English.

## Appendix 1

We studied simulated birdsong to understand how the misclassification of notes affects our inferences about ZLA. We simulated populations in which birdsong either adhered to or did not adhere to ZLA, and we sampled from these populations with and without note classification errors. We studied these samples as described in the body of the paper. We recorded the false positive rate (ie, the proportion of the populations that did not adhere to ZLA but in which we incorrectly inferred ZLA) and the power (ie, the proportion of the populations that adhered to ZLA and in which we correctly inferred ZLA) of our test when notes were correctly or incorrectly classified.

Each population we simulated had a repertoire of *N* note types. We assumed that the expected frequency of use of the note types in the repertoire declined exponentially with the rank order of their frequency of use. In particular, we assumed that the least frequently used note type was used *e^−r^* times as frequently as the most frequently used note type, where *r* was a parameter of the simulation. Thus, the (*i* + 1)^th^ most frequently used note type was used *e^−r/(N−1)^* times as frequently as the *i*^th^ most frequently used note. Each bird in the population had a song composed of *n* notes from this repertoire.

In many real bird populations, the songs of individual birds are not independent because birds learn their songs from others. To simulate this, we allowed each population we simulated to be composed of *L* song lineages (sensu [40]) in which each bird learned his song, possibly with errors, from his father. Each lineage included one founder, all of his sons, all of his paternal grandsons, and so on. For simplicity, we assumed that every lineage included *g* generations after the founder, and that each bird in every generation except the last had *h* sons. Thus, *L*, *g*, and *h* controlled the size of the simulated population. To simulate the song of each founder, we sampled *n* notes, with replacement, from the population repertoire, with each note type chosen with a probability proportional to its expected frequency of use. Thus, each founder’s song might include multiple instances of some note types and no instances of others. To simulate the song of each son, we selected each of the *n* notes in his song independently as follows: with probability *f* we selected a note from his father’s song, with each individual note (not note type) in the father’s song equally likely to be chosen, and we added a note of that type to the son’s song; and with probability (1 − *f*) we selected a note type from the population repertoire, where each note type was chosen with a probability proportional to its expected frequency of use. Thus, the songs of sons were similar but not identical to the songs of their fathers, and the songs of birds from the same lineage were more similar than the songs of birds from different lineages.

To simulate populations in which birdsong adheres to ZLA, we assigned each note type a duration that corresponded to the rank order of its expected frequency of use in the population. Thus, we assigned the shortest duration to the note type with the greatest expected frequency of use, the second shortest duration to the note type with the second greatest expected frequency of use, and so on. To simulate populations in which birdsong does not adhere to ZLA, we assigned each note class a duration independent of its frequency of use.

We simulated three types of classification errors that might occur. For each type of classification error, we simulated *m* misclassifications per population. To simulate the merger of note types, we assumed that note types were most likely to be erroneously merged if their durations were similar. We sampled, without replacement, *m* note types from among the 2^nd^ to *N*^th^ longest note types in the population repertoire, and we merged these note types with the first longer note type that was not among the *m* note types we sampled. For example, if our *m* note types included the 3^rd^ longest note type but not the 4^th^ longest note type then we merged the 2^nd^ and 3^rd^ longest note types, and if our *m* note types included the 3^rd^ and 4^th^ longest note types then we merged the 2^nd^ through 4^th^ longest note types. After mergers, the population repertoire included (*N* − *m*) note types. Because we merged notes with consecutive rank order durations, we could assign rank order durations to note types after mergers without ambiguity. If note classes are erroneously split, the splitting might occur within birds or it might occur among birds. For example, in a real system we might erroneously split within birds if individual birds produce slightly different versions of a note type at different points in their songs, and we might erroneously split among birds if different birds produce slightly different versions of the same note type. For each type of erroneous splitting, we sampled, without replacement, *m* note types from the population repertoire that would be split. To simulate splitting within birds, we considered each individual note in the set of *m* note types that we sampled, and with probability 0.5 we assigned that note to a new note type. To simulate splitting among birds, for each of our *m* note types we considered each bird in the population. If the bird used that note type, then with probability 0.5 we assigned all of that birds’ instances of that note type to a new type. In either case, if note type *i* was among the *m* types we split, all notes that were originally of type *i* but were split from that type were assigned to the same new type*i*′, and after splitting the population repertoires contained (*N* + *m*) note types. We assigned each new note type a duration that fell between the duration of the note type it was created from and the next longest note type in the repertoire.

We simulated two sets of 2 x 10^4^ populations with different parameter values. In each set, we set *L* = 4, *g* = 3, *h* = 2, *r* = 6, *f* = 0.8, and *m* = 10. Thus, each population included 60 birds, the least frequently used note type was used ∼0.0025 times as frequently as the most frequently used note type, and misclassification errors occurred in 10 note types. In the first set of simulations, we set *N* = 60 and *n* = 24. Thus, most birds used only a small subset of note types from the population repertoire. In the second set of simulations, we set *N* = 24 and *n* = 48. Thus, each bird used a larger proportion of the note types available to the population.

Table S1.1 shows the false positive rates and the powers in our simulations with correct note type classification and with each type of classification error. False positive rates highlighted in red are significantly different from 0.05, and powers highlighted in red are significantly different from those with correct note classifications. When note types were erroneously merged, p-values were conservative, and when note types were erroneously split p-values were anticonservative. In some cases, classification errors also reduced the power of the test. When note types were erroneously split among birds, the power of the test appears to increase. This is not surprising, because in this case the test is anticonservative, and thus more likely to indicate ZLA even when there is not relationship between note type duration and frequency of use in the population. It is more surprising that erroneous mergers appear to increase the power in one set of simulations even though the test is conservative. We suspect this is because in this set of simulations each bird used only a small subset of note types, and rare long note types that appeared in the songs of a lineage founder could become disproportionately common in that line. As a result, there was a great deal of stochasticity in the frequencies with which note types were used in the population. Merging note types may help to reduce this stochasticity. When the test for ZLA was well-powered, as in the second set of simulations, tests with correct classification outperformed tests with erroneous mergers.

**Table S1.1.** False positive rates (top) and powers (bottom) in 2 x 10^4^ simulations with correct note type classification and with each type of classification error. False positive rates in red are significantly different from 0.05, and powers in red are significantly different from those with correct note classifications.

| parameter values | correct classification | erroneous mergers | erroneous splitting within birds | erroneous splitting among birds |
| --- | --- | --- | --- | --- |
| $N = 60, n = 24$ | 0.0518<br>0.5203 | **0.0463**<br>**0.5586** | **0.0576**<br>**0.4765** | **0.0574**<br>**0.5400** |
| $N = 24, n = 48$ | 0.0512<br>0.9999 | **0.0430**<br>**0.9906** | **0.733**<br>**0.9886** | **0.0694**<br>1.0000 |

## Appendix 2

Songs archived on Bird-DB are annotated at the level of phrases. Most phrases are monosyllabic – each one consists of a single sound unit (ie, a note) uninterrupted by periods of silence (eg, phrases A and B in figure S2.1). However, some phrase types are polysyllabic (eg, phrases C and D in figure S2.1). These phrases consist of two or more notes separated by short periods of silence. Because these note sequences usually or always appear together in the same order, they were treated as single units in the annotations on Bird-DB [52].

In the body of this paper, we studied Zipf’s law of abbreviation (ZLA) at the level of phrases. Alternatively, one might wish to study ZLA at the level of notes. To this end, we revisited the song annotation for five populations archived on Bird-DB for which exemplars of each phrase type were available. We examined the exemplar for each phrase type, and if sounds within the phrase were separated by clear periods of silence lasting for at least 10 pixels, we divided the phrase type into notes. We classified the individual notes within the phrase classification in Bird-DB phrase as belonging to either the same type (eg, phrase D in figure S2.1) or different types (eg, phrase C in figure S2.1). We observed no cases where notes created by dividing phrases corresponded clearly to existing monosyllabic phrase types, and therefore we treated each note type created by dividing phrases as a new note type in the data. We had only one exemplar per phrase type, so we assumed that every phrase of that type could be divided into the same notes, and that the duration of the new notes would be proportional to their duration in the exemplar.

**Figure S2.1.**
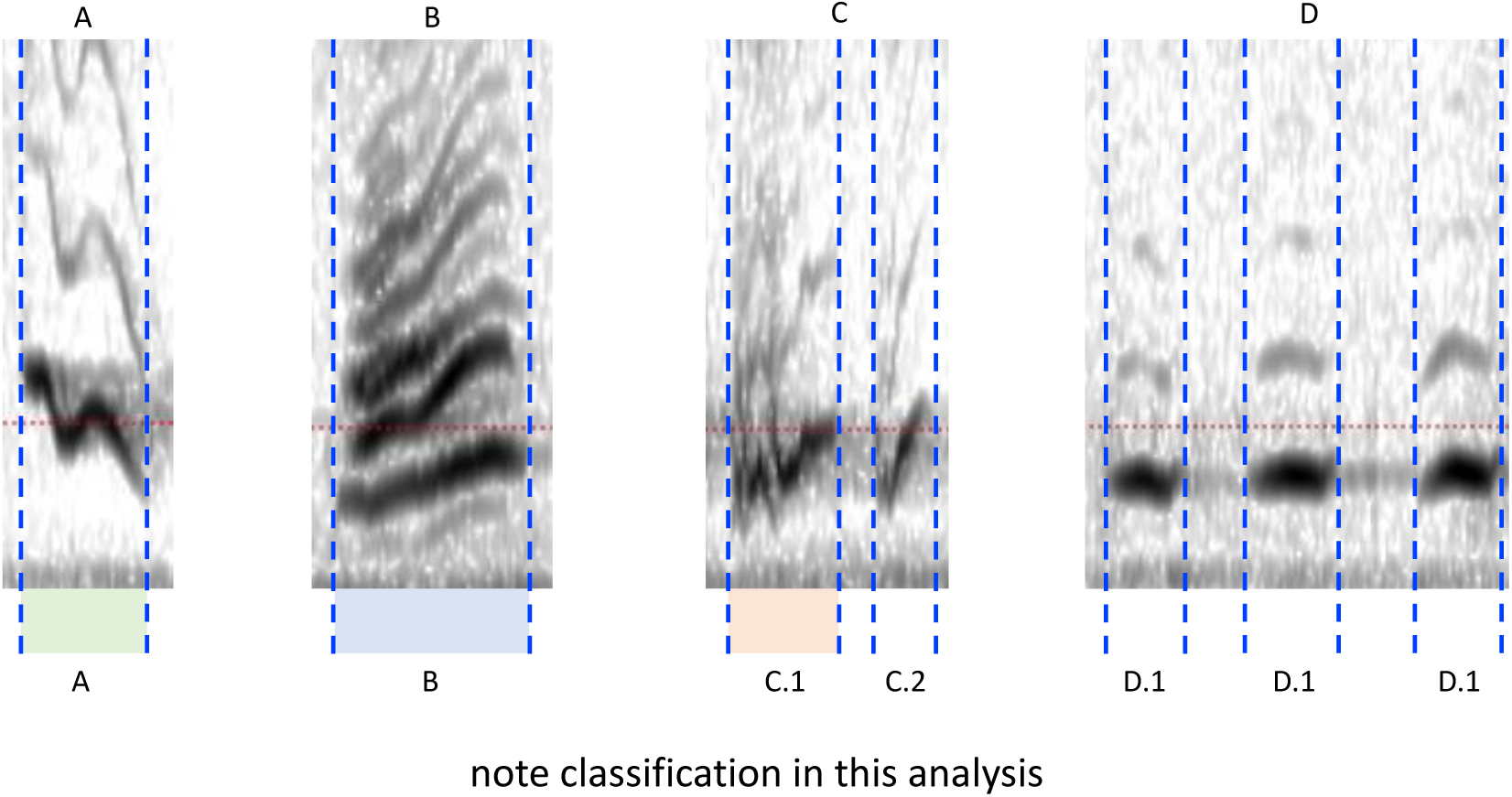
Exemplar phrases from Bird-DB reclassified as notes. Phrases A and B are monosyllabic – each corresponds to a single note type. Phrases C and D are polysyllabic. Phrase C is reclassified as two distinct note types. Phrase D is reclassified as three instances of a single note type. Images are phrases produced by California thrashers and archived on Bird-DB [52].

We repeated the analyses reported in the body of the paper, and we report the results in table S2.1. The results in table S2.1 are qualitatively similar to those reported in the body of the paper. There is little compelling evidence for ZLA in these populations, and if ZLA operates in these populations it operates less strongly than in human languages.

Because we had only one exemplar per phrase type, our re-classification of notes based on exemplars is likely to be less accurate than the classifications at the phrase level provided in Bird-DB, and we do not encourage other researchers to use the re-classification we conducted here. Nonetheless, the results show that qualitative results we obtained by studying ZLA at the population level hold under at least some plausible alternative song annotations.

**Table S2.1.** Summary statistics and concordances in the songs of 5 populations of 3 bird species archived on Bird-DB. Polysyllabic phrases in the original data have been separated into their component notes. Concordances in bold are significantly different from the null expectation (α = 0.10). Statistical tests were one-sided. Thus, p-values less than 0.05 indicate patterns consistent with ZLA.

| species | records studied | total note types | notes record<sup>-1</sup> | note types record<sup>-1</sup> | Shannon diversity | concordance (population) | mean concordance (individual) |
| --- | --- | --- | --- | --- | --- | --- | --- |
| California thrasher | 89 | 809 | 160 | 15.4 | 2.28 | **-0.071 (p = 0.002)** | -0.008 (p = 0.359) |
|  | 7 | 187 | 423 | 59.1 | 3.66 | -0.073 (p = 0.075) | -0.073 (p = 0.063) |
| black-headed grosbeak | 83 | 481 | 166 | 29.1 | 2.98 | 0.002 (p = 0.527) | **-0.051 (p = 0.041)** |
| Cassin's vireo | 13 | 71 | 93.5 | 28.0 | 3.02 | 0.097 (p = 0.878) | 0.015 (p = 0.577) |
|  | 41 | 119 | 101 | 28.3 | 2.92 | -0.007 (p = 0.458) | -0.056 (p = 0.089) |

## Appendix 3

**Figure S3.1.**
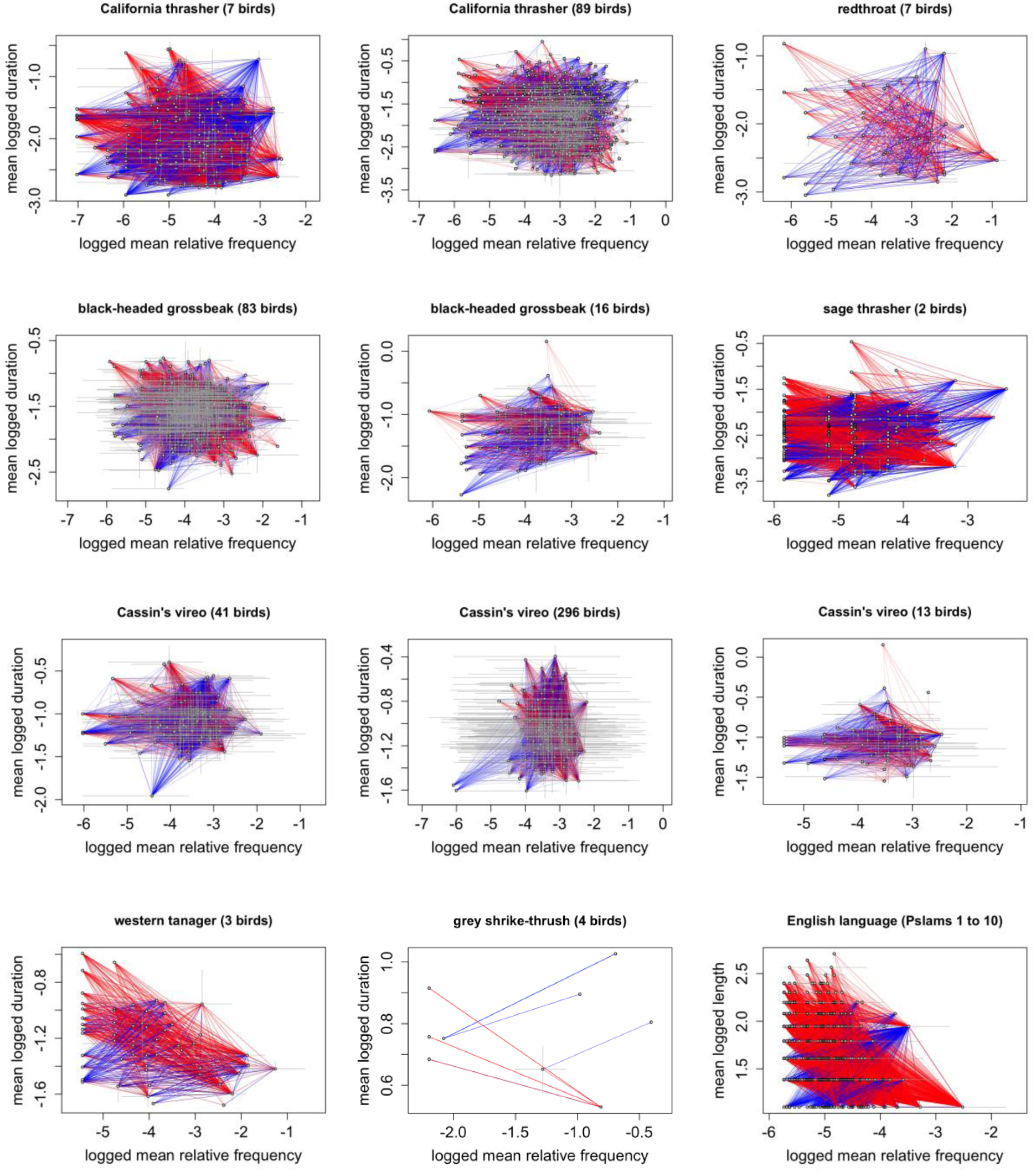
Concordances between note duration and frequency of use in 11 bird populations. Each open circle represents the logged mean frequency of use and the mean logged duration of one phrase type used by the population. Horizontal (vertical) grey lines show the range of logged relative frequencies of use (logged durations) among birds that used the phrase type. Phrase types that appear together in the repertoire of at least one bird are connected by colored lines. Heavier lines indicate that the pair of phrases was used by more birds. Blue (red) lines indicate that the concordance between frequency of use and duration was positive (negative). Intermediate colors indicate that the concordance was positive in some birds and negative in others. For example, this can occur when some birds use the phrase type more frequently than other birds. ZLA predicts that the concordance between frequency of use and duration will be negative. Thus, we should expect to see more red lines in figures when phrase use adheres to ZLA. We found no significant evidence for ZLA in any population except western tanagers (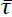 = -0.170, p = 0.038; see table 1). For comparison, the last panel shows the concordance between word length and frequency of use in English based on the first 10 psalms (King James Version), with each psalm treated as if it were produced by a different author. There is strong evidence for ZLA in English based on this sample 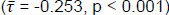.

## Notes

https://doi.org/10.48420/24586791.v1

### Competing Interest Statement

The authors have declared no competing interest.

### Summary of Updates

References updated.

## References

1. Hyland Bruno J, Jarvis ED, Liberman M, Tchernichovski O. Birdsong learning and culture: analogies with human spoken language. Annual Review of Linguistics. 2021;7:449–72. doi: 10.1146/annurev-linguistics-090420-121034.

2. Mori C, Wada K. Songbird: a unique animal model for studying the molecular basis of disorders of vocal development and communication. Exp Anim. 2015;64(3):221–30. doi: 10.1538/expanim.15-0008.

3. Saito N, Maekawa M. Birdsong: the interface with human language. Brain Dev. 1993;15(1):31–9. doi: 10.1016/0387-7604(93)90004-r.

4. Doupe AJ, Kuhl PK. Birdsong and human speech: common themes and mechanisms. Annu Rev Neurosci. 1999;22:567–631. doi: 10.1146/annurev.neuro.22.1.567.

5. Jarvis ED. Evolution of vocal learning and spoken language. Science. 2019;366(6461):50-4. doi: 10.1126/science.aax0287.

6. Lipkind D, Geambaso A, Levelt CC. The development of structured vocalizations in songbirds and humans: a comparative analysis. Topics in Cognitive Science. 2019;12(3):894–909. doi: 10.1111/tops.12414.

7. Bolhuis JJ, Okanoya K, Scharff C. Twitter evolution: converging mechanisms in birdsong and human speech. Nat Rev Neurosci. 2010;11(11):747–59. doi: 10.1038/nrn2931.

8. Fisher SE, Scharff C. FOXP2 as a molecular window into speech and language. Trends in Genetics. 2009;25:166–77. doi: 10.1016/j.tig.2009.03.002.

9. Miller JE, Hafzalla GW, Burkett ZD, Fox CM, White SA. Reduced vocal variability in a zebra finch model of dopamine depletion: implications for Parkinson disease. Physiol Rep. 2015;3(11). doi: 10.14814/phy2.12599.

10. Moorman S, Ahn JR, Kao MH. Plasticity of stereotyped birdsong driven by chronic manipulation of cortical-basal ganglia activity. Curr Biol. 2021;31(12):2619–32 e4. doi: 10.1016/j.cub.2021.04.030.

11. Panaitof SC. A songbird animal model for dissecting the genetic bases of autism spectrum disorder. Dis Markers. 2012;33(5):241–9. doi: 10.3233/DMA-2012-0918.

12. Berwick RC, Okanoya K, Beckers GJL, Bolhuis JJ. Songs to syntax: the linguistics of birdsong. Trends in Cognitive Sciences. 2011;15(3):113–21. doi: 10.1016/j.tics.2011.01.002.

13. Zipf GK. The Psycho-biology of Language: an Introduction to Dynamic Philology. Boston: Houghton Mifflin Company; 1935.

14. Kanwal J, Smith K, Culbertson J, Kirby S. Zipf’s law of abbreviation and the principle of least effort: language users optimise a miniature lexicon for efficient communication. Cognition. 2017;165:45–52. doi: 10.1016/j.cognition.2017.05.001.

15. Bentz C, Ferrer-i-Cancho R. Zipf’s law of abbreviation as a language universal. In: Jäger G, Yanovich I, editors. Proceedings of the Leiden Workshop on Capturing Phylogenetic Algorithms for Linguistics: University of Tübingen; 2016.

16. Linders GM, Louwerse MM. Zipf’s law revisited: spoken dialog, linguistic units, parameters, and the principle of least effort. Psychonomic Bulletin & Review. 2023;30:77–101. doi: 10.3758/s13423-022-02142-9.

17. Ferrer-i-Cancho R, Hernández-Fernández A. The failure of the law of brevity in two New World primates: statistical caveats. Glottotheory. 2013;4(1):45–55. doi: 10.1524/glot.2013.0004.

18. Koshevoy A, Miton H, Morin O. Zipf’s law of abbreviation holds for individual characters across a broad range of writing systems. Cognition. 2023;238:105527. doi: 10.1016/j.cognition.2023.105527.

19. Shu H, Chen X, Anderson RC, Wu N, Xuan Y. Properties of school Chinese: implications for learning to read. Child Development. 2003;74(1):27–47. doi: 10.1111/1467-8624.00519.

20. Kang TS. Linguistic laws and compression in a comparative perspective: a conceptual review and phylogenetic test in mammals. Durham: Durham University; 2021.

21. Semple S, Hsu MJ, Agoramoorthy G. Efficiency of coding in macaque vocal communication. Biology Letters. 2010;6:469–71. doi: 10.1098/rsbl.2009.1062.

22. Clink DJ, Ahmad AH, Klinck H. Brevity is not a universal in animal communication: evidence for compression depends on the unit of analysis in small ape vocalizations. Royal Society Open Science. 2020;7:200151. doi: 10.1098/rsos.200151.

23. Heesen R, Hobaiter C, Ferrer-i-Cancho R, Semple S. Linguistic laws in chimpanzee gestural communication. Proceedings of the Royal Society B - Biological Sciences. 2019;286(1896):20182900. doi: 10.1098/rspb.2018.2900.

24. Huang M, Ma H, Ma C, Garber PA, Fan P. Male gibbon loud morning calls conform to Zipf’s law of brevity and Menzerath’s law: insights into the origin of human language. Animal Behaviour. 2020;160:145–55. doi: 10.1016/j.anbehav.2019.11.017.

25. Bezerra BM, Souto AS, Radford AN, Jones G. Brevity is not always a virtue in primate communication. Biology Letters. 2010;7:23–5. doi: 10.1098/rsbl.2010.0455.

26. Ferrer-i-Cancho R, Lusseau D. Efficient coding in dolphin surface behavioral patterns. Complexity. 2009;14:23–5. doi: 10.1002/cplx.20266.

27. Luo B, Jiang T, Liu Y, Wang J, Lin A, Wei X, et al. Brevity is prevalent in bat short-range communication. Journal of Comparative Physiology A - Neuroethology, Sensory, Neural, and Behavioral Physiology. 2013;199(4):325–33. doi: 10.1007/s00359-013-0793-y.

28. Demartsev V, Gordon N, Barocas A, Bar-Ziv E, Ilany T, Goll Y, et al. The “Law of Brevity” in animal communication: sex-specific signaling optimization is determined by call amplitude rather than duration. Evolution Letters. 2019;3(6):623–34. doi: 10.1002/evl3.147.

29. Hailman JP, Ficken MS, Ficken RW. The ‘chick-a-dee’ calls of Parus atricapillus: a recombinant system of animal communication compared with written English. Semiotica. 1985;56(3-4):191–224. doi: 10.1515/semi.1985.56.3-4.191.

30. Ferrer-i-Cancho R, Bentz C, Seguin C. Optimal coding and the origins of Zipfian laws. Journal of Quantitative Linguistics. 2020;29(2):164–94. doi: 10.1080/09296174.2020.1778387.

31. Ferrer-i-Cancho R, Hernández-Fernández A, Lusseau D, Agoramoorthy G, Hsu MJ, Semple S. Compression as a universal principle of animal behavior. Cognitive Science. 2013;37(8). doi: 10.1111/cogs.12061.

32. Mol C, Chen A, Kager RWJ, ter Haar SM. Prosody in birdsong: a review and perspective. Neuroscience and Biobehavioral Reviews. 2017;81:167–80. doi: 10.1016/j.neubiorev.2017.02.016.

33. Conner RN. Vocalizations of common ravens in Virginia. The Condor. 1985;87:379–88.

34. Favaro L, Gamba M, Cresta E, Fumagalli E, Bandoli F, Pilenga C, et al. Do penguins’ vocal sequences conform to linguistic laws? Biology Letters. 2020;16(2):20190589. doi: 10.1098/rsbl.2019.0589.

35. Lewis RN, Kwong A, Soma M, de Kort SR, Gilman RT. Java sparrow song conforms to Mezerath’s law but not to Zipf’s law of abbreviation. bioRXiv. 2023. doi: 10.1101/2023.12.13.571437.

36. Rehsteiner U, Geisser H, Reyer H-U. Singing and mating success in water pipits: one specific song element makes all the difference. Animal Behaviour. 1998;55:1471–81. doi: 10.1006/anbe.1998.0733.

37. Vallet E, Beme II, Kreutzer M. Two-note syllables in canary songs elicit high levels of sexual display. Anim Behav. 1998;55(2):291–7. doi: 10.1006/anbe.1997.0631.

38. Weiss M, Keifer S, Kipper S. Buzzwords in females’ ears? The use of buzz songs in the communication of nightingales (*Luscinia megarhynchos*). PLoS One. 2012;7(9):e45057. doi: 10.1371/journal.pone.0045057.

39. Suthers RA, Vallet E, Kreutzer M. Bilateral coordination and the motor basis of female preference for sexual signals in canary song. Journal of Experimental Biology. 2012;215(17):2950–9. doi: 10.1242/jeb.071944.

40. Lewis RN, Soma M, de Kort SR, Gilman RT. Like Father Like Son: Cultural and Genetic Contributions to Song Inheritance in an Estrildid Finch. Front Psychol. 2021;12:654198. doi: 10.3389/fpsyg.2021.654198.

41. Lewis RN, Kwong A, Soma M, de Kort SR, Gilman RT. Inheritance of temporal song features in Java sparrows. Animal Behaviour. 2023;206:61–74. doi: 10.1016/j.anbehav.2023.09.012.

42. James LS, Mori C, Wada K, Sakata JT. Phylogeny and mechanisms of shared hierarchical patterns in birdsong. Current Biology. 2021;31(13):2797–808. doi: 10.1016/j.cub.2021.04.015.

43. Fehér O, Wang H, Saar S, Mitra PP, Tchernichovski O. *De novo* establishment of wild-type song culture in the zebra finch. Nature. 2009;459:564–8. doi: 10.1038/nature07994.

44. Greig EI, Taft BN, Pruett-Jones S. Sons learn songs from their social fathers in a cooperatively breeding bird. Proceedings of the Royal Society B - Biological Sciences. 2012;279(1741):3154–60. doi: 10.1098/rspb.2011.2582.

45. Hanley ML. Word Index to James Joyce’s Ulysses. Madison, Wisconsin: University of Wisconsin Press; 1937.

46. Tanimoto AM, Hart PJ, Pack AA, Switzer R. Vocal repertoire and signal characteristics of the ‘alalä, the Hawaiian crow (*Corvus hawaiiensis*). The Wilson Journal of Ornithology. 2017;129(1):25–35. doi: 10.1676/1559-4491-129.1.25.

47. Cohen Y, Nicholson DA, Sanchioni A, Mallaber EK, Skidanova V, Gardner TJ. Automated annotation of birdsong with a neural network that segments spectrograms. eLife. 2022;11:e63853. doi: 10.7554/eLife.63853.

48. Kaewtip K, Alwan A, O’Reilly C, Taylor CE. A robust automatic birdsong phrase classification: A template-based approach. The Journal of the Acoustical Society of America. 2016;140:3691–701. doi: 10.1121/1.4966592.

49. Lachlan RF, Ratmann O, Nowicki S. Cultural conformity generates extremely stable traditions in bird song. Nature Communications. 2018;9:2417. doi: 10.1038/s41467-018-04728-1.

50. Valz PD, McLeod AI. A Simplified Derivation of the Variance of Kendall’s Rank Correlation Coefficient. The American Statistician. 1990;44(1):39–40. doi: 10.1080/00031305.1990.10475691.

51. Youngblood M. Language-like efficiency and structure in house finch song. PsyArXiv. 2023. doi: 10.31234/osf.io/bghqm.

52. Arriaga JG, Cody ML, Vallejo EE, Taylor CE. Bird-DB: a database for annotated bird song sequences. Ecological Informatics. 2015;27:21–5. doi: 10.1016/j.ecoinf.2015.01.007.

53. R Core Team. R: A Language and Environment for Statistical Computing. Vienna, Austria: R Foundation for Statistical Computing; 2023.

54. Ballentine B, Hyman J, Nowicki S. Vocal performance influences female response to male bird song: an experimental test. Behavioral Ecology. 2004;15(1):163–8. doi: 10.1093/beheco/arg090.

55. Podos J, Sung H-C. Vocal performance in songbirds: from mechanisms to evolution. In: Sakata JT, Woolley SC, Fay RR, Popper AN, editors. The Neuroethology of Birdsong. Springer Handbook of Auditory Research: Springer Cham; 2020. p. 245–68.

56. Hsu MJ, Chen L-M, Agoramoorthy G. The vocal repertoire of Formosan macaques, *Macaca cyclopis*: acoustic structure and behavioral context. Zoological Studies. 2005;44(2):275–94.

57. McLachlan JR, Magrath RD. Speedy revelations: how alarm calls can convey rapid, reliable information about urgent danger. Proceedings of the Royal Society B - Biological Sciences. 2020;287:20192772. doi: 10.1098/rspb.2019.2772.

58. Suzuki TN. Parental alarm calls warn nestlings about different predatory threats. Current Biology. 2011;21(1):R15–R6. doi: 10.1016/j.cub.2010.11.027.

59. Suzuki TN. Communication about predator type by a bird using discrete, graded and combinatorial variation in alarm calls. Animal Behaviour. 2014;87:59–65. doi: 10.1016/j.anbehav.2013.10.009.

60. Tanimoto AM, Hart PJ, Pack AA, Switzer R, Banko PC, Ball DA, et al. Changes in vocal repertoire of the Hawaiian crow, *Corvus hawaiiensis*, from past wild to current captive populations. Animal Behaviour. 2017;123:427–32. doi: 10.1016/j.anbehav.2016.11.017.

